# A synthetic phageome reveals widespread and diverse human antibody responses to bacteriophages

**DOI:** 10.64898/2026.10.29.762592

**Authors:** Katarzyna Gembara, Marek Harhala, Paulina Miernikiewicz, Zuzanna Kaźmierczak, Paul Bollyky, Andrzej Konieczny, Aleksandra Wilczak, Izabela Rybicka, Piotr Tynecki, Mirosława Różycka, Natalia Jędruchniewicz, Marlena Kłak, Daniel Nelson, Wojciech Witkiewicz, Krystyna Dąbrowska

**Author notes:** equal (Katarzyna Gembara, Marek Harhala).

## Abstract

Bacteriophages colonize humans, yet their recognition by humoral immunity remains unclear. We developed PhageScan, a PhIP-Seq platform comprising 277,051 phage and 49,630 human viral oligopeptides. Profiling sera from 192 healthy donors without prior phage therapy characterized anti-phage response specificity and revealed widespread, strong IgG reactivity to phages, comparable in prevalence and magnitude to that against human viruses. Recognition patterns were however highly individualized. Among phage sequences, 12,576 peptides were significantly enriched in at least two individuals. Strongly immunogenic epitopes preferentially originated from phages infecting pathogenic bacteria, whereas widely shared “public” epitopes were enriched among phages of environmental and food-associated bacteria. ImmE, a lactococcal phage endolysin recognized in 59.9% of individuals, showed reduced antibacterial activity in the presence of sera identified as ImmE-reactive by PhageScan, demonstrating functional relevance. These findings establish phage-directed antibodies as a widespread component of human immunity and PhageScan as a scalable platform for defining phage–antibody interactions.

---

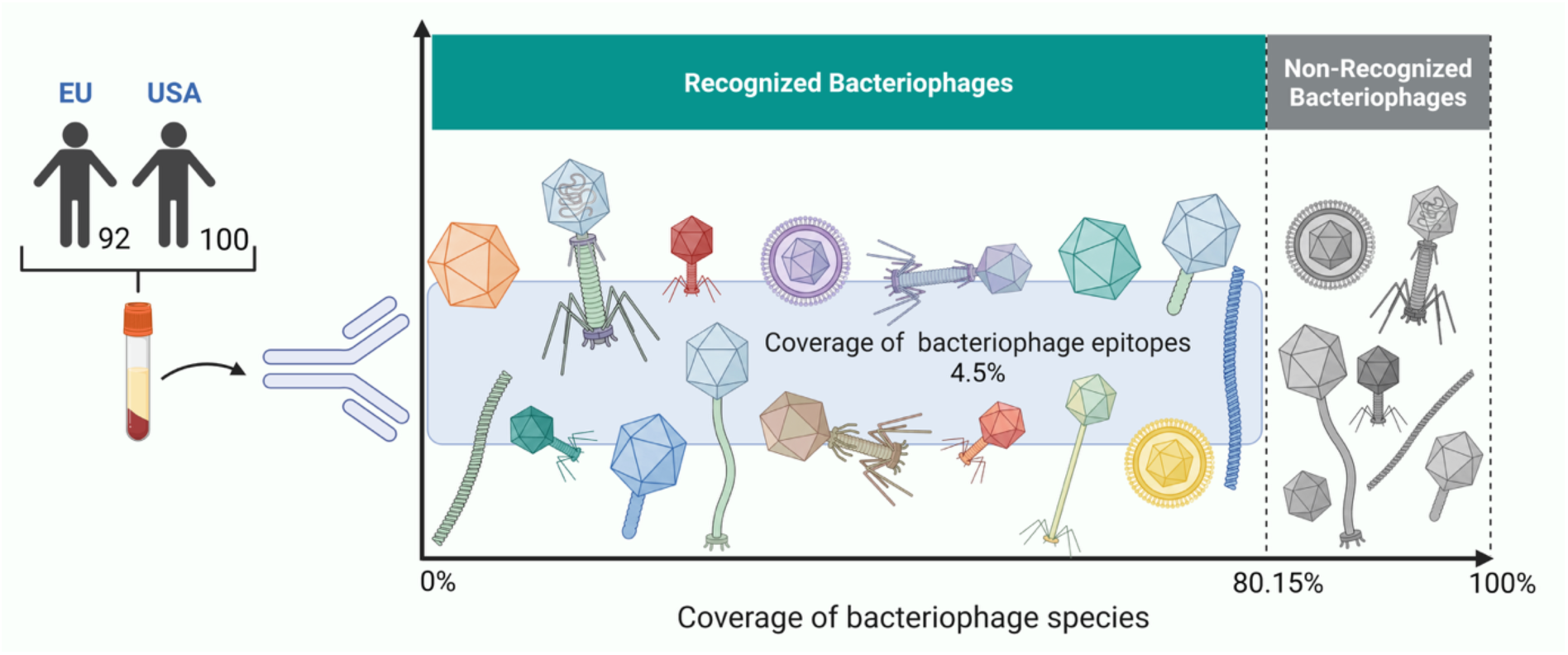

## Introduction

Bacteriophages, viruses that parasitize bacteria, form complex, diverse communities naturally present in human microbiomes. They can treat bacterial infections, but even without therapeutic administration, our tissues are continuously exposed to bacteriophages generated at bacterial colonization sites that then circulate throughout the body (1–4). Each person has a unique history of natural contact with phages (5–7).

Studies of lytic phages, including those used therapeutically against bacterial infections, clearly demonstrate that phages are immunogenic and elicit antibody responses (8–10). However, much of this anti-phage immunity is directed against endogenous phages; even patients who have never received phage therapy often have pre-existing antibodies that can recognize phages, including those used in therapeutic preparations (11–14). This can lead to faster and more efficient neutralization of phages and even dramatic reduction in their antibacterial activity (15,16). In the context of phage therapy, such specific neutralization may result in poor pharmacokinetics and reduced lytic phage therapy efficacy (17).

Available data are limited to antibodies against one or a few phages. The prevalence, magnitude, diversity, and composition of phage-directed antibody responses in human populations remain unclear. How these responses compare to immunity against pathogenic human viruses and which phage epitopes are typically recognized at the molecular level also remain largely unknown.

Phage ImmunoPrecipitation Sequencing (PhIP-Seq) enables population serological profiling (18). This approach enables the detection and relative quantification of human serum IgG and identifie tens of thousands of immunogenic epitopes. It is based on libraries of short oligopeptides displayed on T7 bacteriophage capsids (phage display), which interact with IgG antibodies present in serum samples. Peptides that do not bind antibodies are removed, allowing a comprehensive and precise characterization of those recognized by human IgG (19). In recent years, the method has advanced rapidly, with the development of improved protocols, dedicated software tools, and several important review studies (20–22). PhiP-Seq has been applied to serological profiling of human populations, particularly for human viruses, where it is known as VirScan (23).

Here, we provide a first insight into human serological profiles against circulating bacteriophages by introducing PhageScan, a PhIP-Seq approach incorporating a structural phage pan-proteome (UniProt) including 277,051 phage-derived oligopeptides as well as 49,630 oligopeptides derived from human viruses. We compare phage-specific responses with those against human viruses and identify phage epitopes commonly recognized by human IgG.. We further investigate the functional implications of PhageScan findings and how they translate into the neutralizing activity of human IgGs.

## Materials and methods

### Study participants and serum samples

Serum samples were obtained from 192 healthy adult donors, including 92 individuals from Poland and 100 from the United States. Blood was collected into clotting tubes and allowed to clot for 0.5-1 h at room temperature. Serum was separated by centrifugation at 2,000×g for 15 min at room temperature and stored at −20°C until analysis.

### Ethics statement

This study was approved by the Bioethics Committee of the Regional Specialist Hospital in Wrocław (approval no. KB/02/2017).

Human serum samples from the Polish participants were collected within the project, each participant received detailed information about the study and provided written informed consent before sample collection. Human serum samples from the US participants were obtained commercially from BioIVT (Westbury, NY, USA). Samples were provided in deidentified form, with only donor age, sex, and race available to the investigators, and no information permitting identification of individual donors was provided. Accordingly, use of these specimens did not constitute human subjects research under the U.S. Department of Health and Human Services regulations for the protection of human subjects (45 CFR 46.102[e]), and IRB approval was not required.

### PhageScan library design and serological profiling

PhageScan was adapted from the VirScan method developed by Xu et al. (23) to identify phage-derived oligopeptides recognized by human IgG. All bacteriophage genomes are available in GenBank (National Center for Biotechnology Information. They were accessed 1 June 2018 and were screened for annotated open reading frames (ORFs). ORFs containing the terms “structural,” “tail,” “decoration,” “capsid,” “scaffold,” “baseplate,” “coat,” “virion,” “structure,” “lysin,” “muramidase,” or “holin” were selected, whereas those containing “initiator,” “chaperone,” “maturation protease,” or “morphogenesis” were excluded.

The selected protein sequences were computationally divided into 56-amino-acid oligopeptides, starting every 28 amino acids and thus providing 50% overlap to retain linear epitopes spanning fragment boundaries. Following deduplication, the sequences were reverse-translated using codons optimized for expression in *Escherichia coli*. The oligonucleotide library was synthesized using SurePrint nucleotide-printing technology (Agilent) and cloned into a phage-display system using the T7Select 415-1 Cloning Kit (Merck Millipore), with Liga5™ (A&A Biotechnology) replacing T4 DNA ligase and the standard EcoRI and HindIII restriction sites.

Because large-scale serological profiling of phage-derived sequences had not previously been validated, 49,630 oligopeptides derived from human viruses and previously used in VirScan were included as a positive reference. The phage-derived fraction comprised 277,051 oligopeptides, resulting in a combined library of 326,681 oligopeptides.

### Immunoprecipitation

Immunoprecipitation of the library was performed in accordance with a previously published protocol (23). Briefly, the phage library was amplified in a standard culture as described in the manufacturer’s manual, then purified using hollow fiber filtration membrane. In the final step, the purified and concentrated library was suspended in Phage Extraction Buffer (20 mM Tris-HCl, 100 mM NaCl, 60 mM MgSO_4_, pH 8.0). All plastic containers (96-well plates) used for immunoprecipitation were prepared by blocking with 3% Bovine Serum Albumin (BSA) in TBST buffer overnight on a rotator (40 rpm, 4°C). A sample representing an average of 2x10^5^ copies of each clone in 250 µL was mixed with 1 µL of human serum (two technical replicates were applied) and incubated overnight at 4°C with rotation (50 rpm). A 20 μL aliquot of a 1:1 mixture of Protein A and Protein G Dynabeads (Invitrogen) was added and incubated for 4 h at 4°C with rotation (50 rpm). Liquid in all wells was separated from Dynabeads on a magnetic stand and removed. Beads were washed 5 times with 280 µL of a wash buffer (50 mM Tris-HCl, pH 7.5, 150 mM NaCl, 0.1% Tween-20) and beads were resuspended in 60 µL of water to elute the immunoprecipitated bacteriophages from the beads.

The immunoprecipitated part of the library was then used for amplification of the insert region according to the manufacturer’s instructions with a Phusion Blood Direct PCR Kit (Thermo Fisher Scientific). The primers T7_Endo_Lib_LONG_FOR GCCCTCTGTGTGAATTCT and T7_Endo_Lib_LONG_REV GTCACCGACACAAGCTTA were used and a second round of PCR was carried out with the IDT for Illumina UD indexes (Illumina Corp.) to add adapter tags. Sequencing of the amplicons in accordance with Illumina next generation sequencing (NGS) technology was outsourced (Genomed, Warszawa). Sequencing data generated in this study have been deposited under project accession number PRJEB124668.

### Sequencing data analysis

Sequenced amplicons along with 23 control samples (*input sample*), a phage library before immunoprecipitation, were mapped to the original nucleotide library sequences by the bowtie2 software as published by Xu et al. (23). NGS sequencing reads were mapped by the bowtie2 software package with use of the file with list of oligonucleotide sequences synthesised originally to create a library as indexes (end-to-end mode “-q - 5 9 --no-unal --no-hd --no-sq --ignore-quals --mp 3 --rdg 150,100 --rfg 150,100 --score-min L,-0.6,-0.6”). The number of hits that mapped to each reference sequence (highest score) was counted (count, c).

The signal in each sample was calculated according to a formula (1).

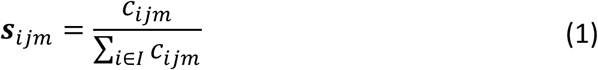

s – signal of i-th sequence in j-th serum sample and m-th technical replicate

c – number of reads mapped to i-th sequence in m-th technical replicate of j-th serum sample

I – set of all reference sequences (used as indexes to in mapping by the bowtie2 software)

Signals from oligopeptides not detected in a given technical replicate were treated as left-censored and assigned the minimum signal observed in that replicate. This conservative approach reduced the risk of classifying oligopeptides as false positives. We calculated log_10_(**s**), mean log10(**s**) and its standard deviation from all *input samples* for each sequence separately. Next, we compared each log10(**s**) in samples to mean and standard deviation of a respective sequence from *input samples* and calculated a chance for this value to be accidental (normal distribution). Relative signal (enrichment): an average signal in technical replicates of the sample divided by the average signal in *input samples* of the same sequence is a signal ratio (relative signal). Sequences with relative signal below cut-off probability in both technical replicates were marked as “detected” in that serum sample.

Phage-derived oligopeptides were classified as immunogenic when they were significantly enriched (p-value<0.00001 in both experimental replicates) in sera from at least two individuals. This minimum threshold was applied to reduce the contribution of random detection events. Public epitopes were defined as oligopeptides significantly enriched in at least 20% of the analyzed sera. Top-immunogenic oligopeptides were defined as sequences significantly enriched in at least two sera at p<10^−14^. Subsequent enrichment analyses included four analytical sets: (i) the reference library, comprising oligopeptides detected in the physical input library before immunoprecipitation; (ii) immunogenic oligopeptides; (iii) public epitopes; and (iv) top-immunogenic oligopeptides. The counts of oligopeptides in each set were compared using chi-squared test for independence with Yates correction.

### Data extraction

Using the “olgname” column from the source file we extracted 32,276 unique protein accession numbers. Next, Entrez API from BioPython v1.81 package (24) was adopted to download amino acid sequences with their meta-data (id, description, source, organism, db_source, data_file_division, molecule_type and date). We calculated hash checksums (SHA-256) (25,26) for each amino acid sequence to be sure that we have unique samples in the repository.

From the “db_source” column we extracted 5,401 unique bacteriophage accession numbers. Once again Entrez API from BioPython package were used to download phage genome sequences with their meta-data (accession number, host, lab_host, islolation_source, name, strandedness, moltype, topology, division, dates, taxonomy and comment). Again, we calculated the hash checksums (SHA-256) for each genome sequence to validate that 5,061 of the total 5,401 are unique. The rest of the phage genome were removed as duplicates with different accession numbers. “host”, “lab_host” and “islolation_source” columns were normalized by domain experts. When applicable, they were assigned to super-groups defining phage types. In case of data unavailability or ambiguity, the relevant hit was excluded from analysis. A detailed description of the curation and categorization of the GenBank data for phage host and isolation source is provided in the Supplementary Table S1.

### PhageScanMut library design and identification of residues contributing to antibody recognition

Public epitopes, defined as immunogenic oligopeptides significantly enriched in at least 20% of the tested sera, were subjected to systematic alanine-substitution scanning using an approach adapted from Xu et al. (23). For each public epitope, single residues or windows of two or three consecutive residues were replaced with alanine. Native alanine residues were replaced with glycine. This procedure generated up to 56 single-, 55 double-, and 54 triple-substitution variants per epitope, corresponding to a maximum of 165 mutant oligopeptides for each parental sequence. Each mutant differed from the parental sequence only at the designated substituted position or window.

The resulting PhageScanMut library was synthesized, cloned, displayed, immunoprecipitated, sequenced, and analyzed using the same experimental and computational workflow as the original PhageScan library. For each mutant oligopeptide, the number of sera in which it was significantly enriched was recorded. A significant reduction in the number of recognizing sera relative to other variants of the same parental oligopeptide was interpreted as evidence that the substituted residue or region contributed to antibody recognition.

To identify substituted residues statistically, the median number of positive sera was calculated for each parental (non-substituted) oligopeptide, each substitution type separately. For a given parental epitope, the null probability of recognition was defined as the median number of positive sera divided by the total number of sera tested. The probability of observing the number of positive sera obtained for each mutant was then calculated using a lower-tailed binomial test, with the total number of tested sera as the number of trials. The resulting p-values were adjusted for multiple testing using the Benjamini–Hochberg procedure, with a false discovery rate of 5%. Substitutions with adjusted P values below 0.05 were classified as moderately affecting antibody recognition, whereas those with adjusted p-values below 0.0005 were classified as strongly affecting antibody recognition. For double- and triple-substitution variants, the classification was assigned to the first residue of the substituted window.

### Phage species burden score

Because phage species are represented unequally in public sequence databases, and consequently in the PhageScan library, raw numbers of recognized oligopeptides cannot be compared directly across species. Extensively studied phages were represented by larger numbers of proteins and oligopeptides and therefore have a greater probability of recognition, including through cross-reactive antibodies. To account for this representation bias, we calculated a burden score comparing the observed contribution of each phage species to all detected oligopeptide-recognition events with its corresponding contribution to the original library:

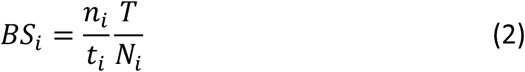

BS_i_ – burden score of *i-th* phage species

n_i_ – the number of oligopeptides representing the *i-th* phage species in all tested sera

t – total number of immunogenic oligopeptides identified in all sera

N_i_ – the number of oligopeptides representing the *i-th* phage species in original library of oligopeptides

T – total number of oligopeptides representing all phage species in the original library of oligopeptides

For calculations of p-value we used binomial model with probability equal to ratio of the number of sequences representing i-th phage species to all sequences in original library (Ni/T).

The sum of all detected sequences representing i-th phage species and sum of all detected sequences in the presented research were used in binomial model as the number of tested events and total number of events. P-values calculated by SciPy package and Python Programming Language (25,26). P-values adjusted for multiple hypotheses with were calculated using Benjamini-Hochberg method (false discovery rate) in statsmodel package (‘fdr_bh’) (27,28).

Species-specific detection thresholds were derived using an approach adapted from Xu et al. (23) to account for differences in the number of proteins representing individual species in the library. Briefly, linear regression was performed between the log-transformed mean signal and the log-transformed number of unique proteins represented in the library for each species. The fitted regression slope was used to scale the detection threshold for each phage species, using a threshold of three recognized peptides for herpes simplex virus 1 (HSV1) as the reference. A serum sample was classified as seropositive for a given phage species—and thus as showing evidence of previous exposure—when the number of significantly enriched oligopeptides assigned to that species was equal to or greater than its species-specific threshold (29).

### Epitope homologies

For consistency and clarity, each oligopeptide was annotated using one arbitrarily selected representative phage record. Sequence homology between recognized phage-derived oligopeptides and eukaryotic organisms was assessed using BLAST searches against the complete database. Homologies for selected public epitopes were additionally identified in NCBI (accessed 10 March 2026).

A total of 12,576 oligopeptide sequences (immunogenic oligopeptides) were screened for homology against the eukaryotic subset of the NCBI non-redundant protein sequence database (NR), restricted to Eukaryota (taxid: 2759) and Unclassified Eukaryota (taxid: 42452). Searches were performed using DIAMOND in blastp mode with the very-sensitive preset. A maximum of five hits was reported per query, and a permissive E-value threshold of 10 was used during the initial search.

Following the database search, individual hits for all 12,576 sequences were subjected to quality filtering. Entire query sequences were not excluded. Instead, individual hits were removed when the matched organism or database entry belonged to one of the following categories: (i) bacterial organisms, including *Escherichia coli*, Bacteria (unresolved), *Pseudomonas aeruginosa*, *Streptococcus pyogenes*, *Vibrio lentus*, *Sphingomonas* spp., *Sphingobium yanoikuyae*, *Pseudomonas soli*, *Stenotrophomonas* spp., *Escherichia* spp., *Kocuria* spp., Enterobacteriaceae, the *Enterobacter cloacae* complex, *Klebsiella michiganensis*, *Achromobacter marplatensis*, *Salmonella enterica* subsp. enterica serovar Newport, *Enterococcus avium*, *Cedecea neteri*, and *Paracidovorax* spp.; (ii) bacteriophage-related or vector entries, including *Tequatrovirus* T4 (bacteriophage T4) and the lambda gt10 cloning vector; and (iii) entries with non-specific annotations, namely “synthetic construct” and “unidentified.” These hits were excluded because they represented prokaryotic or artificial entries within the eukaryotic search space or lacked sufficient taxonomic resolution for meaningful biological interpretation.

For each query sequence from which one or more hits were removed, the remaining valid hits were shifted to consecutive positions while preserving their original ranking. Any remaining hit positions were left blank. Overall, at least one hit was removed from 204 query sequences. Sequence significance was then recalculated using only the retained hits. A sequence was classified as significant when at least one retained hit had an E-value ≤ 0.05 and as not significant otherwise.

All 12,576 query sequences were retained for downstream analysis. Of these, 1,359 (10.81%) were classified as significant, whereas 11,217 (89.19%) were classified as not significant. All subsequent analyses, including hit-count distributions, E-value and sequence-identity distributions, query-coverage statistics, and organism-level summaries, were performed using the complete filtered dataset.

### Machine learning–based prediction of phage lifestyle and protein function

For bacteriophage life cycle prediction we used PhageAI API v0.4.2 (30) . That two-cascade AI model is able to predict if a bacteriophage is chronic, temperate or virulent with a high probability. To make classification the complete genome phage sequence is required only. Next, PhageAI API was also executed to classify phage structural proteins. Selected method is classifying amino acid sequence as structural protein (Baseplate, Collar, Head tail, Major capsid, Major tail, Minor capsid, Minor tail, Portal, Tail fiber, Tail sheath) or as AA sequence with other functions.

All the predictions data (classes and probabilities) from PhageAI API were included in the final source file for further analysis.

### ImmE protein expression, isolation, and purification

A synthetic ImmE coding sequence was cloned into the pBAD/HisA expression vector using the restriction enzymes NcoI and BglII; a C-terminal His tag was attached to ImmE via a triple-serine linker (BioCat GmbH, Heidelberg, Germany). Recombinant ImmE was expressed in *Escherichia coli* TOP10 cells (Thermo Fisher Scientific) cultured in LB medium supplemented with ampicillin (100 µg/mL). Cultures were grown at 37 °C to an OD₆₀₀ of 1.0–1.2. Protein expression was induced with 0.25% (w/v) L-arabinose, followed by incubation at 30 °C for 4 h.

Cells were harvested by centrifugation and resuspended in phosphate buffer (50 mM NaH₂PO₄, 300 mM NaCl, pH 7.5) containing lysozyme (1.5 mg/mL) and phenylmethylsulfonyl fluoride (PMSF; 1 mM). The suspension was incubated on ice for 6–7 h and then frozen overnight at −80 °C. After thawing, DNase (up to 30 µg/mL) and RNase (up to 60 µg/mL) were added together with Mg²⁺ (up to 1 mM). Insoluble material was removed by two successive centrifugation steps at 16,000 × g for 30 min at 4 °C.

The clarified supernatant was sequentially passed through 0.45- and 0.22-µm sterile filters, supplemented with imidazole to a final concentration of 10 mM, and incubated with Ni-NTA agarose. After binding, the resin was washed, and ImmE was eluted stepwise with phosphate buffer containing increasing concentrations of imidazole (25–500 mM). Fractions containing ImmE were pooled, dialyzed against PBS, and subjected to endotoxin removal using EndoTrap® HD resin (LIONEX GmbH, Braunschweig, Germany). The final protein preparation was dialyzed against PBS, sterile-filtered, and quantified spectrophotometrically.

## Fluorometric assay

The bacteriolytic activity of ImmE in the presence of 10% human serum was evaluated using a SYTOX Green–based fluorometric assay adapted from a previously described method (31). Three *Lactococcus lactis* strains from the Polish Collection of Microorganisms (HIIET; PCM473, PCM2379, and PCM2853) were used as target bacteria. Overnight cultures grown in brain heart infusion (BHI) medium (Oxoid) were inoculated into fresh BHI medium and incubated at 37 °C until the late logarithmic growth phase. Bacterial cells were collected by centrifugation at 7,000 × g for 7 min at 4 °C, washed twice with PBS, and resuspended in PBS to an OD₆₀₀ of 0.6.

Assay mixtures were prepared in PBS and contained SYTOX Green nucleic acid stain, bacterial cells, ImmE (1 µg/mL), and/or human serum, according to the experimental condition. Control samples included the dye, serum, protein, or bacteria alone, as well as the corresponding combinations. The final reaction volume was 200 µL per well, and each condition was tested in duplicate. Fluorescence kinetics were measured at room temperature in black 96-well plates using a BioTek microplate reader with excitation and emission wavelengths of 485 and 528 nm, respectively.

Raw fluorescence signals (RFU) were first corrected for background fluorescence and auto-fluorescence by subtracting a matrix-matched background for each specific reaction environment. For every time point (t), the fluorescence of the control sample lacking endolysin was subtracted from the corresponding enzyme-treated sample within the same environment:

For the buffer condition (PBS):

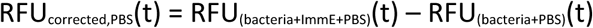

For each individual serum condition:

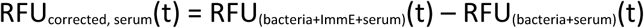

To standardize initial baseline differences across the various sera and the PBS control, ΔRFU relative to t = 0 was calculated as follows:

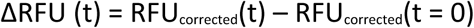

### Statistics and data processing

Statistical analyses were performed using Statistica (StatSoft Polska Sp. z o.o., Kraków, Poland) and in Python 3. Chi-square tests between all combination of presented groups were performed using GraphPad Prism version 10.2.0 for Windows, GraphPad Software, Boston, Massachusetts USA. Odds ratios (ORs) and 95% confidence intervals (CIs) were calculated in R (version 4.3.2) using RStudio (version 2023.12.1). Statistical significance was assessed using two-sided Fisher’s exact tests, with P < 0.05 considered significant. Spearman’s rank correlation coefficients (ρ) and the corresponding two-sided P values were calculated in R using the cor.test function with method = "spearman".

In Statistica software, data preprocessing included division into training and test sets, one-hot encoding of categorical variables, and class balancing using undersampling, oversampling and the Synthetic Minority Oversampling Technique (SMOTE). Weight-of-evidence-based categorization and predictor ranking were also performed. Univariable and multivariable logistic regression, including stepwise models, was used to assess associations between phage and bacterial host characteristics and immunogenicity or public-epitope status. Models were fitted with and without phage-family classification. Associations were expressed as odds ratios with 95% confidence intervals, and regression coefficients were evaluated using Wald tests, with P < 0.05 indicating statistical significance. Logistic model discrimination was assessed using receiver operating characteristic curves and the corresponding area under the curve (ROC-AUC) in the training and test sets. Associations between categorical variables were examined using Pearson’s and likelihood-ratio chi-square tests. Complementary classification analyses used CART and XGBoost models, evaluated using accuracy, sensitivity and specificity. XGBoost performance was additionally assessed using ROC-AUC and F1-score, with a decision threshold of 0.6. Exploratory k-means clustering evaluated solutions with two, three and four clusters; associations between cluster membership and discordant immunogenicity classifications were assessed using chi-square tests.

### Use of AI tools

Machine learning was used for prediction of phage lifestyle and protein function (see above). ChatGPT powered by GPT-5.6 (OpenAI; accessed August 2026) (32) was used for English-language editing and stylistic refinement. All AI-assisted text was critically reviewed and edited by the authors, who take full responsibility for the final content. No other aspects of writing included AI.

## Results

### Specific IgG responses to bacterial viruses resemble those directed against human viruses

To characterize human IgG responses to bacteriophages, we performed high-throughput serological profiling of healthy individuals from two geographically distinct populations, Poland and the United States, using PhageScan a synthetic phageome library (Fig. 1). The library represented the complete phage proteome available in GenBank and comprised 277,051 phage-derived oligopeptides originated from structural proteins and bacteriolytic non-structural proteins, including endolysins, holins, and depolymerases. A set of 49,630 oligopeptides derived from human viruses, based on VirScan (23), served as a positive reference. Both fractions were analyzed together in one pool, enabling direct comparison of IgG responses to phages and human-infecting viruses.

**Figure 1.**
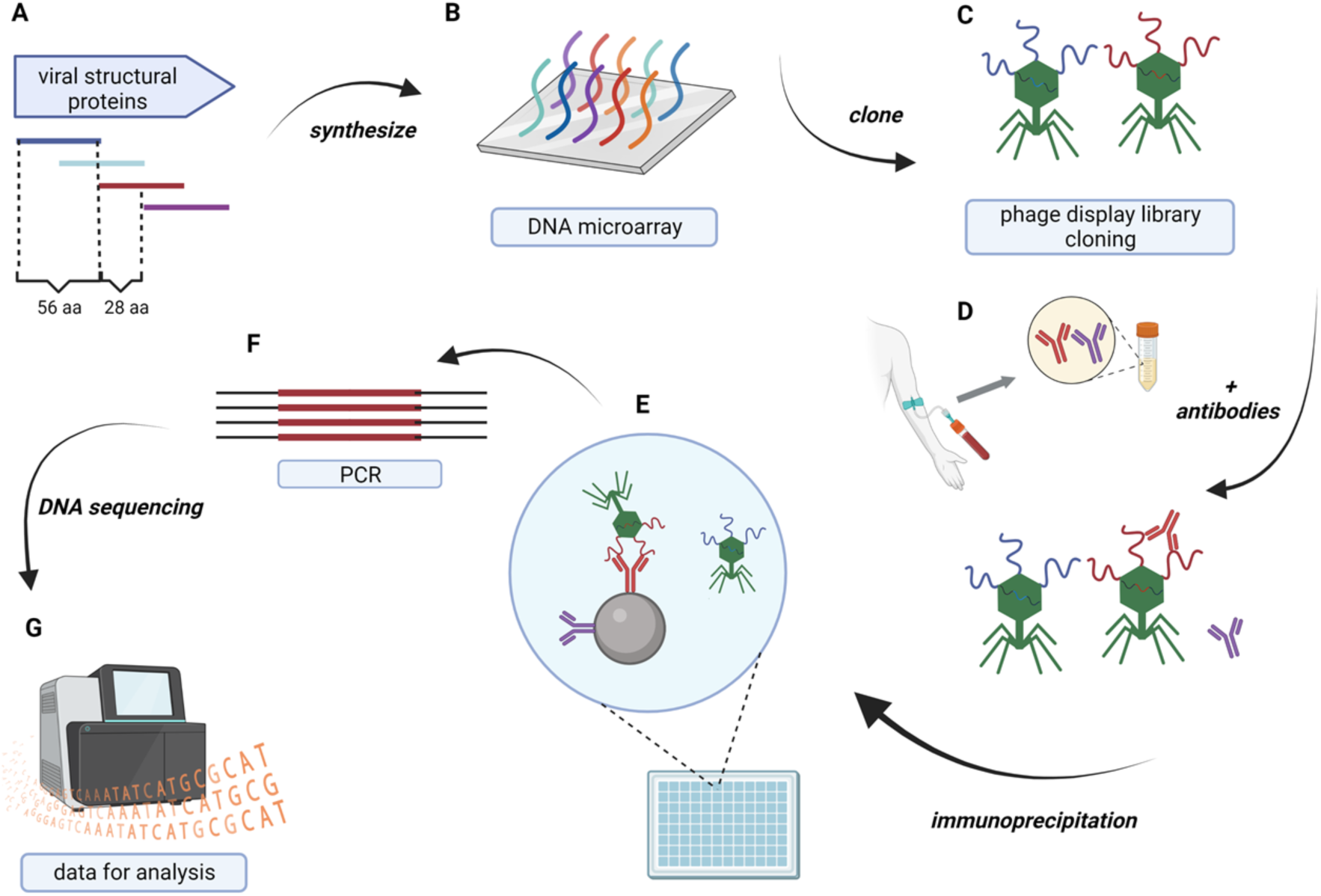
PhageScan technology. PhageScan, a phage-display serological profiling platform to identify bacteriophage-derived peptides recognized by human IgG included following steps: **A.** Bacteriophage genomes in GenBank were screened for annotated open reading frames encoding proteins likely to be exposed to the immune system, including structural, capsid, tail, coat, baseplate, lysin, and holin proteins. **B.** Oligopeptides were synthesized for these selected proteins after dividing them into 56-amino-acid peptides with 28-amino-acid offsets, creating 50% overlap to preserve linear epitopes spanning peptide boundaries. C. These oligopeptides were cloned into a T7 phage-display system. The resulting library contained 277,051 phage-derived peptides plus 49,630 human viral peptides from VirScan as positive controls, for 326,681 total peptides. **D.** For serological profiling, approximately 2 × 10^5^ copies of each clone were incubated with human serum in duplicate overnight. **E.** Antibody-bound phages were captured using Protein A/G magnetic beads, extensively washed, and eluted. **F.** DNA inserts **G.** from immunoprecipitated phages were amplified by PCR, indexed with Illumina adapters, and analyzed by next-generation sequencing. This approach enables high-throughput identification of phage-derived peptide sequences targeted by circulating human IgG antibodies. (adapted from VirScan, (23) (BioRender).

For each recognized oligopeptide, the median relative signal was plotted against its recognition frequency across the study population. The overall IgG-response profile for phage-derived oligopeptides resembled that for human viruses, including a similar range of response strengths (Fig. 2A,B). Recognition frequency distribution were also similar. Phage-derived oligopeptides tended to be recognized in more individuals than those derived from human viruses, although not statistically significant.

**Figure 2.**
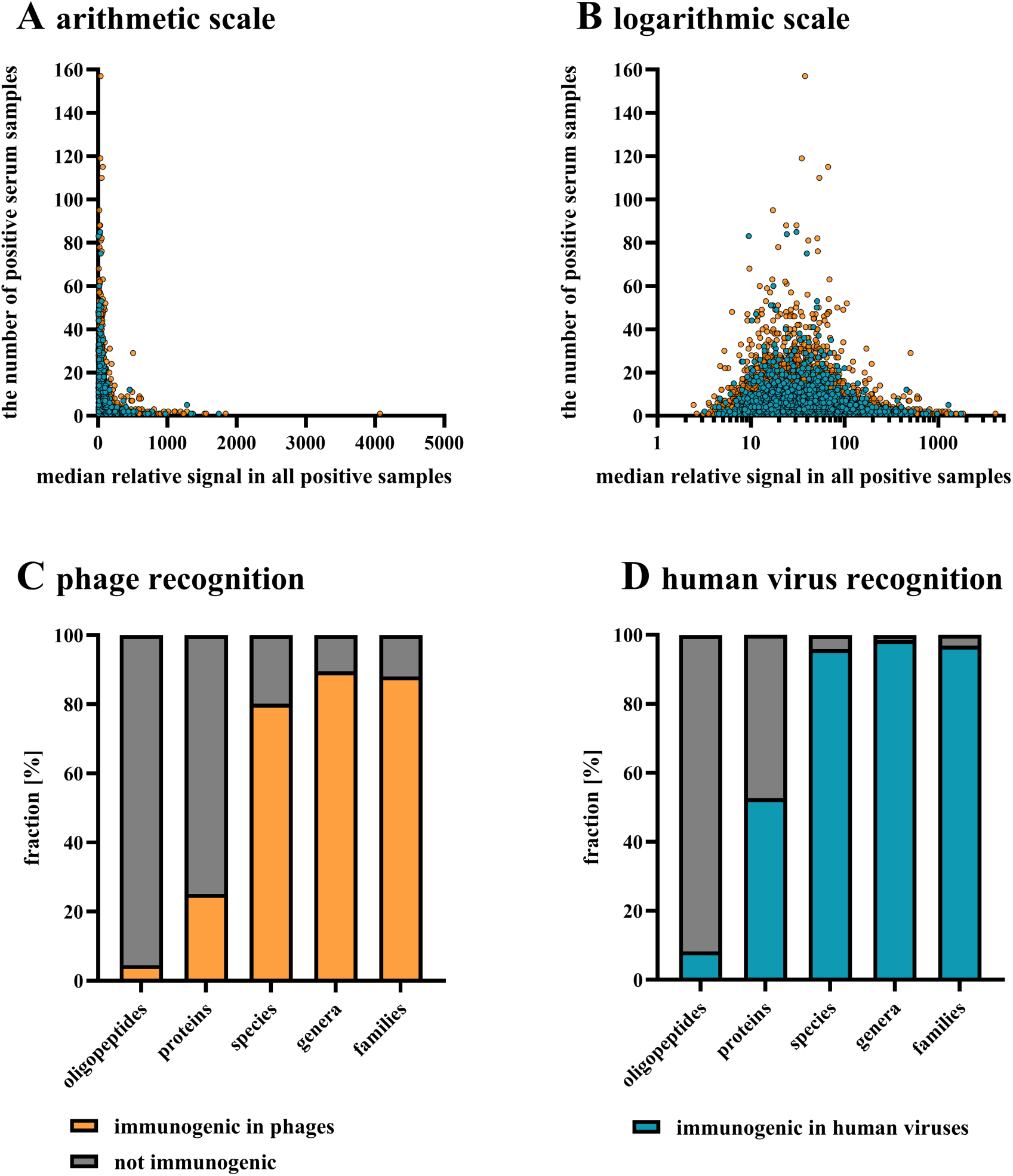
Comparison of human IgG responses to phage- and human virus-derived oligopeptides. Recognition frequency across the study population and median response strength for phage-derived (orange) and human virus-derived (blue) oligopeptides are shown on linear (**A**) and logarithmic (**B**) scales. IgG recognition coverage at the oligopeptide, protein, species, genus, and family levels is shown for phages (**C**) and human viruses (**D**).

The burden score is a library-normalized measure of virus-specific IgG recognition that corrects for differences in virus representation within the oligopeptide library. It is a normalized measure of antibody recognition, not a direct measure of viral load, infection prevalence, or disease burden (29). In the combined ranking, phages and human viruses were interspersed, further demonstrating comparable overall patterns of IgG recognition across both viral groups (Supplementary Table S2).

For both phages and human-infecting viruses, IgG antibodies recognized only a small fraction of the represented oligopeptides and proteins, whereas recognized sequences were distributed across a large proportion of represented viral taxa (Fig. 2C,D). Nevertheless, recognition of the phageome was less extensive than that of the human virome: IgGs recognized 4.5% of phageome oliopeptides versus 8.2% of viral oligopeptides and 25.2% of phageome proteins versus 52.7% of viral proteins, respectively. At the taxonomic level, recognized sequences represented approximately 80–90% of phage species, genera, and families, compared with more than 95% of the corresponding taxa among human-infecting viruses.

### Identification and classification of IgG-reactive phage-derived oligopeptides

Following immunoprecipitation with human sera, phage clones displaying 12,576 phage-derived oligopeptides (4.5% of the library) showed significant enrichment in samples from at least two individuals. These oligopeptides were therefore classified as immunogenic. Among them, 87 were classified as public epitopes because they were significantly recognized by IgG in at least 20% of the individuals analyzed. In addition, 637 oligopeptides met the criteria for the top-immunogenic set, showing highly significant enrichment following immunoprecipitation with sera from at least two individuals (p<10^−14^) (Fig. 3A). The complete list of identified oligopeptides is provided in Supplementary Table S3.

**Figure 3.**
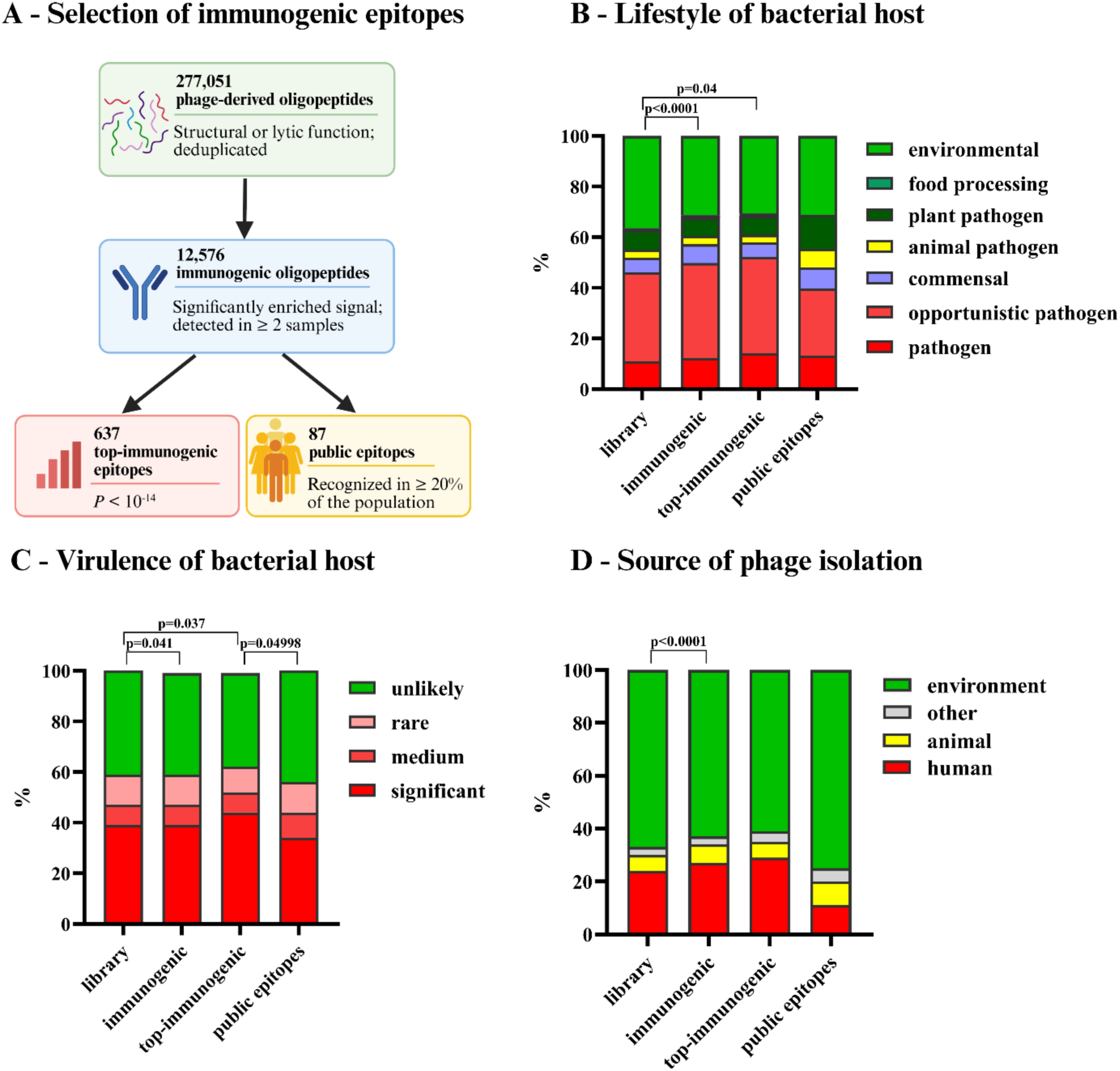
Biological characterization of phage-derived oligopeptides recognized by human IgG antibodies. **A.** Identification pipeline for immunogenic, highly immunogenic, and public epitopes; **B.** Immunogenicity of phages analyzed according to the lifestyle of their bacterial hosts; **C.** Immunogenicity of phages analyzed according to the virulence of their bacterial hosts; **D.** Immunogenicity of phages analyzed according to the source of phage isolation.

Subsequent analyses compared the reference library with three sets of recognized oligopeptides: immunogenic, public, and top-immunogenic.

### Public and top-immunogenic epitopes derive from phages infecting different bacterial host groups

We used curated GenBank metadata on phage isolation source and bacterial host to characterize the phages represented by immunogenic oligopeptides. Phage-derived oligopeptides were grouped according to bacterial host and isolation source, and their representation in the immunogenic, public, and top-immunogenic sets was compared with that in the reference library (Fig. 3B–D). Although the differences were modest, two contrasting patterns emerged.

Highly immunogenic epitopes (those eliciting the strongest IgG responses) were more frequently derived from phages infecting pathogenic bacteria. In contrast, public epitopes were more frequently associated with phages infecting environmental bacteria, including bacteria found in plant-derived foods.

Oligopeptides from phages infecting pathogenic bacteria accounted for 52% of the top-immunogenic set, compared with 39% of the public set and 46% of the reference library (Fig. 3B). Similarly, oligopeptides from phages infecting bacteria capable of causing human infection represented 44% of the top-immunogenic set, compared with 34% of the public set (P < 0.05) and 39% of the reference library (Fig. 3C).

These host-associated patterns were mirrored by phage isolation sources, which were classified as environmental (e.g., water, soil, and plants) or human-associated (e.g., human biological samples and urban or hospital sewage). Phages isolated from environmental sources contributed more frequently to public epitopes, whereas those from human-associated sources contributed more frequently to the top-immunogenic set (Fig. 3D). Consistent with these observations, univariable logistic regression showed that oligopeptides from phages infecting environmental rather than pathogenic bacteria had lower odds of belonging to the top-immunogenic set (OR = 0.755, 95% CI: 0.743–0.767, P < 0.0001). The odds were also lower for phages infecting bacteria unlikely to cause human infection than for those infecting bacteria posing a substantial threat to human health (OR = 0.791, 95% CI: 0.770– 0.815, P < 0.001).

Phage lifestyle showed limited association with immunogenicity. Univariable logistic regression revealed no difference between virulent and temperate tailed phages. Phages with a chronic infection cycle, however, were underrepresented in the immunogenic set relative to virulent phages (OR = 0.569, 95% CI: 0.436–0.744, P < 0.001; data not shown).

Phage morphology was not associated with the odds of an epitope belonging to the public or top-immunogenic sets. In the broader immunogenic set, however, podovirus-like oligopeptides were modestly enriched relative to all other morphotypes combined, including myovirus-like, siphovirus-like, filamentous, and tailless icosahedral phages (OR = 1.079, 95% CI: 1.022–1.138, P < 0.01; data not shown).

Overall immunogenicity did not differ significantly among structural protein classes. However, tail-derived oligopeptides were modestly more frequent in the top-immunogenic set than in the public set (χ² test, P = 0.037).

### Identification of amino acids critical for phage antigen interaction with human IgGs in public epitopes

Public epitopes and selected sequences from the model bacteriophage λ (101 oligopeptides) were analyzed to identify individual amino acid residues critical for sequence-specific IgG binding. A new library was designed and synthesized, with substitutions at each position. Most substitutions were to alanine, while alanine residues were replaced with glycine. To capture potential local structural effects, we included single-, double-, and triple-site mutations. This library was used for immunoprecipitation with the same human sera. Loss of enrichment for an oligopeptide variant indicated that the substituted residue was critical for IgG binding to the epitope. This approach enabled detailed molecular characterization of public epitopes and identified the key amino acid determinants underlying their recognition by human IgG antibodies (Figure 4, full list in Supplementary Material, File 1).

**Figure 4.**
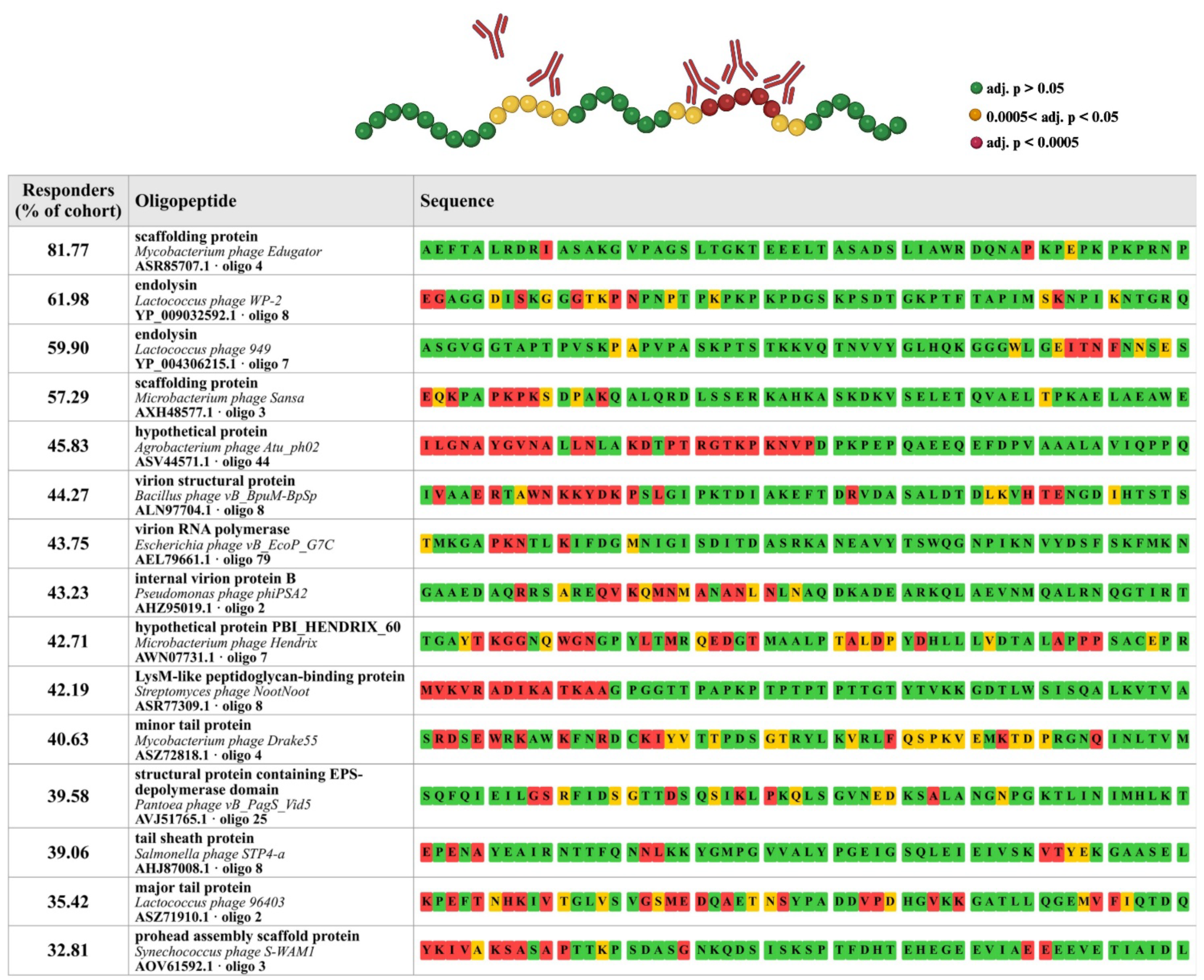
Amino acid residues critical for human IgG recognition in the 15 most frequently recognized phage-derived oligopeptides. Critical amino acid residues were identified by immunoprecipitation of an amino acid substitution library. A complete overview of the residues identified in public epitopes and the model bacteriophage λ is provided in Supplementary File 1.

### Homologies between phage, bacteria, and eukaryotic antigens

Most recognized oligopeptide sequences occurred in multiple phages, commonly with sequence identities approaching 100%. Representative homologies of ≥95% for selected public epitopes are presented in Supplementary Table S4. Phages sharing highly similar proteins typically belonged to closely related biological groups with similar origins and host ranges.

High sequence identity was also observed between phage-derived oligopeptides and bacterial sequences (Supplementary Table S4).

To evaluate potential cross-reactivity with antibodies induced by unrelated, non-phage antigens, we analyzed the homology of recognized phage-derived oligopeptides to eukaryotic sequences. Over 6% of oligopeptides showed significant homology, whereas almost 94% did not (Figure 5A). Some oligopeptides displayed high sequence identity to eukaryotic proteins (Figure 5B) and/or numerous matches within a single eukaryotic species (Figure 5C). The most frequent matches involved fungi, protists, and plants. A complete analysis is provided in Supplementary File 2.

**Figure 5.**
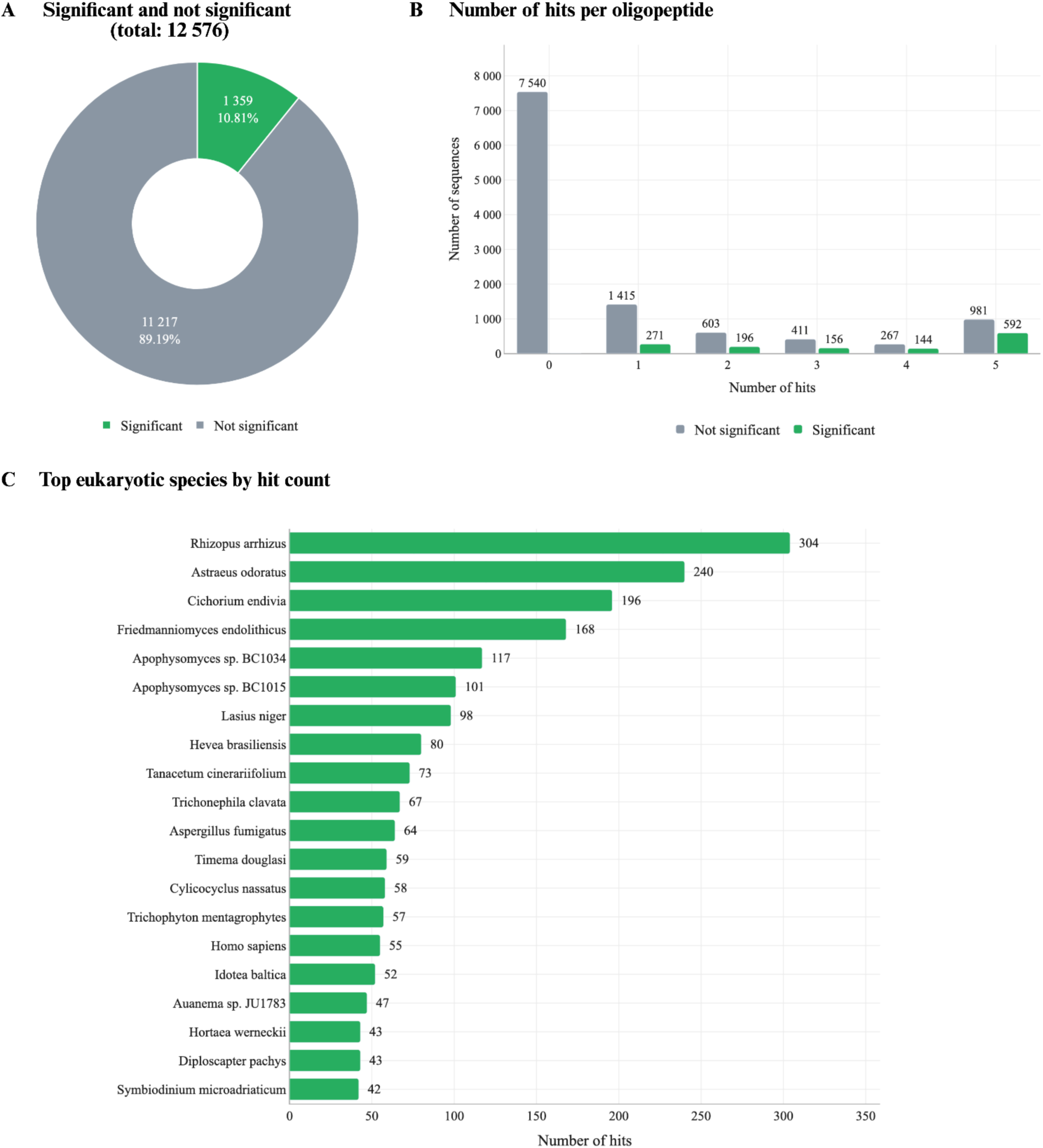
Sequence homology between IgG-recognized phage-derived oligopeptides and eukaryotic proteins. **A.** Proportions of recognized phage-derived oligopeptides with and without significant homology to eukaryotic sequences. **B.** Sequence identity distributions between phage-derived oligopeptides and their eukaryotic homologues. **C.** Number of homologous matches for the 20 eukaryotic species with the highest numbers of hits. A full list is available in Supplementary File 2.

### Geographically distinct populations show similar patterns of phage recognition

Despite differences between the Polish and US populations in the recognition of individual phage-derived oligopeptides, with 2,649 and 2,765 oligopeptides recognized exclusively in each group, respectively (Supplementary Table S5), their overall IgG-recognition profiles were highly similar. No significant population-level differences were detected in the distribution of recognized oligopeptides according to bacterial host category, including environmental, pathogenic, commensal, and other bacteria (Pearson’s χ^2^, p=0.09056; likelihood-ratio χ^2^, p=0.09028) Profiles also did not differ by the medical relevance of the bacterial host (p=0.090415 for both tests), phage isolation source (p=0.66710 and p=0.66670), protein category (p=0.55266 and p=0.55170), phage morphology/family (p=0.48214 and p=0.48221), or life cycle (p=0.65210 and p=0.63754, for Pearson’s and likelihood-ratio χ^2^ tests, respectively) (Fig. 6). The same oligopeptide ranked highest in both populations. Thus, despite differences in individual oligopeptide recognition, broader patterns of phage-directed IgG recognition were consistent across the two geographically distinct populations (Supplementary Table S5).

**Figure 6.**
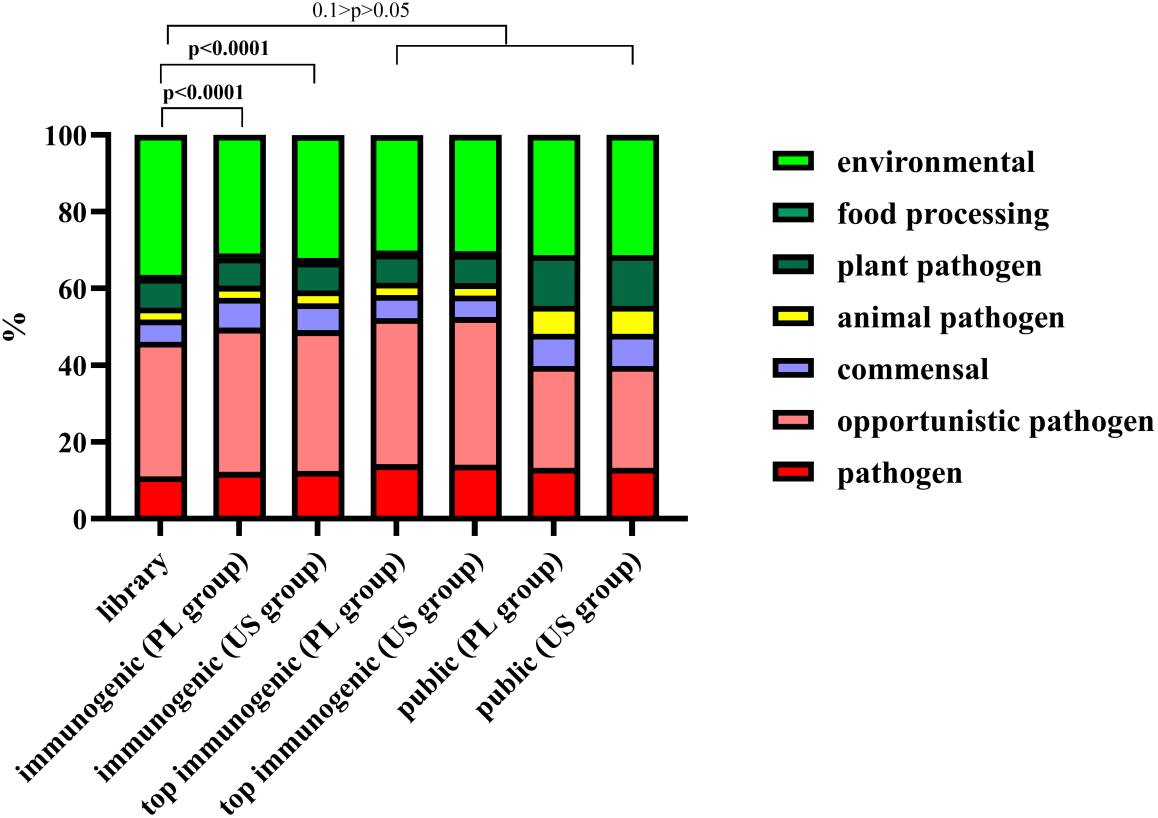
Distribution of immunogenic oligopeptides across United States of America (US) and Polish (PL) populations in relation to phage host biology. No statistical significant differences were observed between US and PL groups by Chi-square tests.

### PhageScan signals correlate with ELISA reactivity and identify sera that inhibit a phage endolysin

To determine whether PhageScan detected antibody recognition was associated with functional activity, we focused on a widely recognized antigen: an endolysin from a Lactococcus phage (NCBI accession YP_004306215.1) (33), termed ImmE (immunogenic endolysin). ImmE-specific IgG was detected in 59.9% of sera. The public epitope was mapped onto the AlphaFold-predicted three-dimensional structure of ImmE (Fig. 7A). It lay outside the enzymatically active domain (EAD), in a separate domain for which InterProScan identified no known function; this domain may be involved in bacterial cell wall binding.

**Figure 7.**
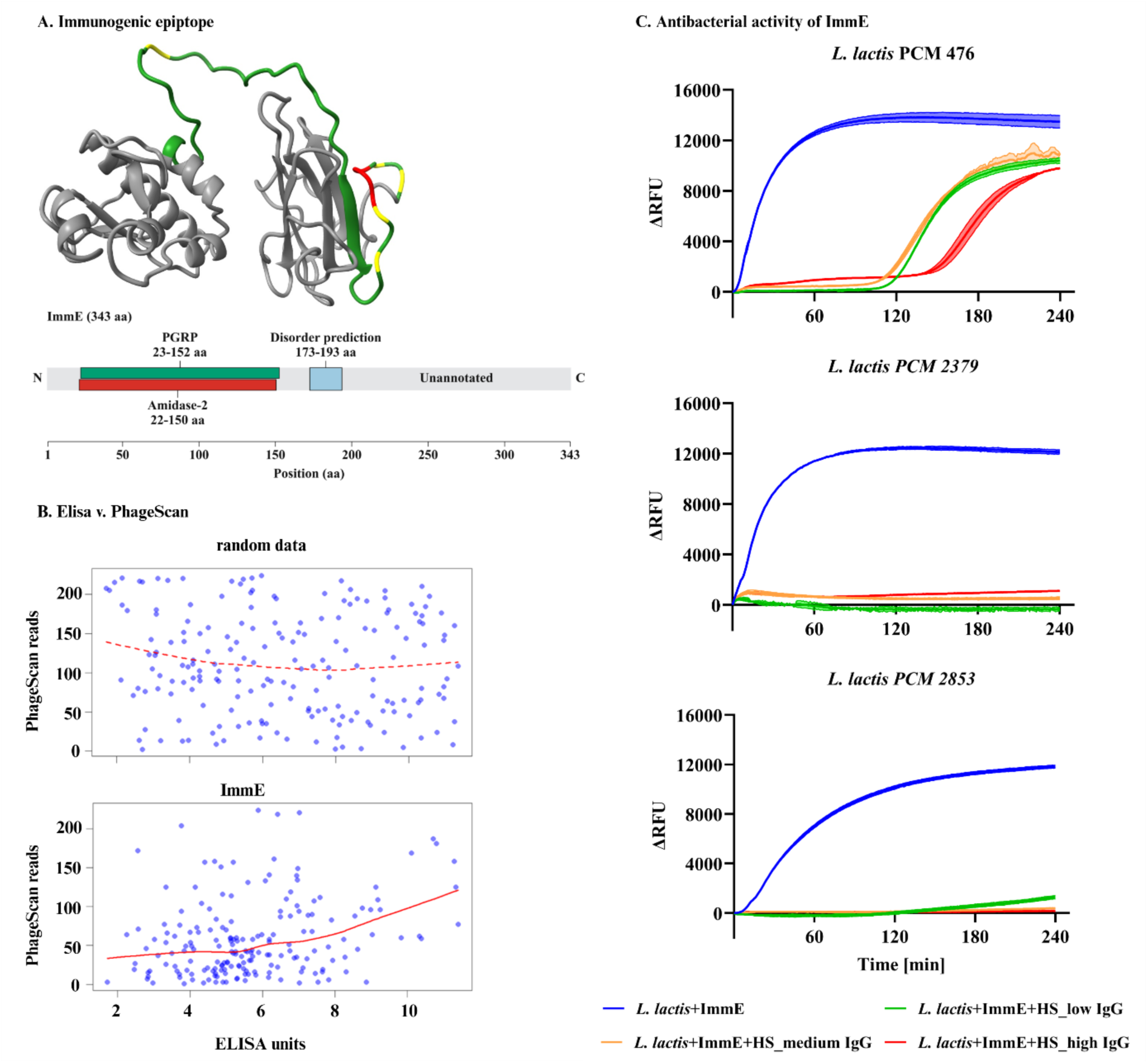
Structural, serological, and functional validation of a public ImmE endolysin epitope identified by PhageScan. **A.** Location of the public epitope within the AlphaFold-predicted three-dimensional structure of ImmE. Green - low IgG, yellow - medium IgG, red - high IgG. **B.** Spearman correlation between ELISA units and PhageScan reads - NGS read counts normalized as mean relative signal, both reflecting serum recognition of ImmE epitopes. The corresponding correlation obtained using randomly generated control values is shown for comparison. **C.** Inhibition of ImmE bacteriolytic activity by selected anti-ImmE IgG-positive human sera against three *Lactococcus lactis* strains. *L. lactis* + ImmE (blue), activity of ImmE alone; *L. lactis* + ImmE + HS_low (green), ImmE activity in the presence of serum with low anti-ImmE reactivity (PhageScan signal: 10); L. lactis + ImmE + HS_medium (orange), ImmE activity in the presence of serum with medium anti-ImmE reactivity (PhageScan signal: 77); and *L. lactis* + ImmE + HS_high (red), ImmE activity in the presence of serum with high anti-ImmE reactivity (PhageScan signal: 187). ImmE was used at 1 µg/mL, and serum-containing reactions included 10% (v/v) human serum. Bacteriolytic activity was measured using a SYTOX Green-based fluorometric assay adapted from a previously described method (31). ΔRFU, change in relative fluorescence units at each indicated time point relative to the beginning of the measurement. Higher ΔRFU values indicate stronger antibacterial activity of ImmE.

We first compared the normalized number of PhageScan NGS reads (PhageScan signal) mapping to the ImmE epitope with normalized ELISA reactivity (ELISA units; Supplementary Table S6). These measurements showed a modest but statistically significant positive correlation (Spearman’s ρ = 0.245, P < 0.001; Fig. 7B), indicating that the PhageScan signal reflects antigen-specific IgG reactivity measured by ELISA.

We next selected three representative sera with high, intermediate, and low normalized PhageScan signals for the ImmE epitope (187, 77, and 10 reads, respectively) and assessed their effects on ImmE bacteriolytic activity using a SYTOX-based fluorometric assay (PMID: 34759903). ImmE alone showed marked bacteriolytic activity against all three *Lactococcus lactis* strains, whereas this activity was reduced with the tested sera. All three sera inhibited ImmE activity, with the overall inhibition broadly associated with the PhageScan signal, although its magnitude and kinetics varied among strains (Fig. 7C).

## Discussion

Here, we show that antibody recognition of bacteriophages is a widespread feature of human immunity, detected in every individual examined despite there being no history of phage therapy. IgG reactivity spanned a broad taxonomic range of phages and a highly diverse set of phage proteins, indicating that phages encountered as components of the human phageome or through food, water, and other environmental exposures are not immunologically silent. Although phages do not replicate in human cells, the population-level profile of IgG responses to phage-derived oligopeptides resembled that observed for human viruses, showing pronounced interindividual variation and a comparable breadth and frequency of epitope recognition.

Despite this broad reactivity, phage-specific antibody profiles were highly individualized. Of the 277,051 phage-derived oligopeptides examined, 12,576 (4.5%) showed significant serum reactivity, yet only 87 (0.03%) were recognized by antibodies in at least 20% of individuals (Fig. 4A and Supplementary Table S2 and S3). Analyzing the geographically distinct US and Polish cohorts separately increased the number of commonly (≥20%) recognized oligopeptides only modestly, to 108 and 118, respectively. By contrast, more than 2,500 oligopeptide reactivities were characteristic of each cohort (Supplementary Table S5), indicating substantial population-associated differences in epitope recognition. Thus, reduced within-cohort heterogeneity in geographic, ethnic, dietary, and lifestyle factors had little effect on the scarcity of public epitopes, although these factors may shape which specific epitopes are recognized.

Consistent with this recognition heterogeneity, stratification of immunogenic oligopeptides by the biological characteristics of their source phages revealed only limited trends. One such trend was that the most immunogenic oligopeptides were more frequently derived from phages infecting pathogenic bacteria than from phages associated with environmental bacteria. Because phage ecology is closely linked to that of their bacterial hosts, exposure to phages associated with bacterial pathogens may accompany human bacterial infections (34,35). Such infections elicit strong innate immune responses that shape the subsequent development of adaptive immunity (36). We therefore hypothesize that the inflammatory environment and increased antigen exposure associated with bacterial infection may also promote the recognition of pathogen-associated phages, resulting in stronger phage-specific IgG reactivity than that elicited by phages encountered primarily in environmental settings. Future studies linking individual infection histories with phage-specific antibody profiles will help determine the contribution of bacterial infections to these responses. This question is particularly relevant to phage therapy, as therapeutic phages are administered during active infections and therefore encounter an infection-activated immune environment.

The potential impact of pre-existing immunity on phage therapy (11,37) prompted us to examine epitopes recognized across a substantial proportion of the population. Such public epitopes may identify phages that are commonly targeted by human antibodies and whose antibacterial efficacy could therefore be reduced in vivo. Using site-directed mutagenesis, we mapped the amino acid residues essential for IgG recognition within these epitopes (23) (Fig. 3; complete list in Supplementary File 1). This resource may support the development of therapeutic phage collections, particularly off-the-shelf preparations, for which the probability of encountering patients with pre-existing antibodies is especially relevant.

Unexpectedly, public epitopes were not restricted to structural phage proteins but were also identified in enzymes such as endolysins and depolymerases (Fig. 3). Antibody recognition of these proteins may be functionally important because enzyme-specific antibodies can inhibit their antibacterial activity (38). We therefore investigated ImmE, an endolysin derived from a lactococcal phage (33) and recognized by IgG in 59.9% of the examined sera (Figs. 3 and 7A). ImmE showed strong antibacterial activity against three *Lactococcus lactis* strains, whereas ImmE-reactive serum samples inhibited this activity. These findings provide functional validation of the epitopes identified by PhageScan and demonstrate that common antibody recognition can directly affect the antibacterial activity of a phage-derived protein.

To our knowledge, this is also the first experimental indication that naturally occurring human antibodies may protect potentially beneficial bacteria, such as lactococci, by neutralizing a phage-encoded lytic enzyme. Extracellular phage-mediated lysis can occur without productive infection, as exemplified by lysis from without, while endolysins released during phage-induced lysis may potentially damage neighboring susceptible cells. Antibodies targeting phages or their lytic enzymes could therefore limit such effects and help preserve beneficial bacterial populations. This possibility warrants further investigation, as it suggests that anti-phage antibodies may not only interfere with phage therapy but may also have a beneficial role in protecting commensal or probiotic bacteria.

Although PhageScan reactivity, as verified for ImmE, correlated significantly with ELISA measurements, the correlation was only low to moderate (Fig. 7B). This limited concordance likely reflects, at least partly, differences in antigen format and the types of epitopes captured by the two assays. PhIP-Seq primarily detects linear epitopes, whereas ELISA using intact proteins may also detect conformational epitopes formed by residues that are distant in the primary sequence (39). ELISA, however, measures overall antibody binding to a protein but does not identify the recognized epitopes or directly predict the functional consequences of binding. By contrast, PhageScan maps IgG-reactive oligopeptides to specific proteins and their domains or, more broadly, to defined components of phage particles. This resolution is important because the general term “anti-phage antibodies” does not distinguish between antibodies targeting different phage components or their potentially distinct functional effects (40).

The extensive sharing of homologous sequences and functional modules among bacteriophages (41,42) has important implications for interpreting individual immunogenic epitopes. Because many proteins and oligopeptides are shared by multiple phages, antibodies elicited by one phage may cross-react with homologous sequences in others. Moreover, each oligopeptide in our study was annotated using a single, arbitrarily selected representative phage record from NCBI. Detection of antibodies against a given oligopeptide therefore neither identifies the antigen or phage that elicited the response nor limits its potential functional relevance to the annotated phage (Supplementary Table S4). Matches between immunogenic phage-derived oligopeptides and bacterial genomes were also common (Supplementary Table S4) and may represent integrated temperate phages, defective prophages, or sequences transferred between phages and their hosts.

Among the 12,576 immunogenic phage-derived oligopeptides, a subset also showed significant sequence similarity to eukaryotic proteins (Fig. 6). These similarities raise the possibility that antibodies elicited by environmental, dietary, or host-associated eukaryotic antigens-including proteins from fungi such as *Rhizopus*, food plants such as chicory, and various insects, arachnids, nematodes, crustaceans, and even human-may cross-react with phage-derived oligopeptides. However, sequence similarity alone does not demonstrate immunological cross-reactivity. Some matches may also result from contamination or misannotation of deposited eukaryotic genome assemblies with microbial or phage-derived sequences. These findings should therefore be interpreted cautiously and experimentally validated before cross-reactivity is inferred.

ASR85707.1_oligopeptide_4 may illustrate such cross-reactivity. It was the most frequently recognized oligopeptide in our cohort (81.8%) and was assigned to a scaffolding protein of mycobacteriophage Edugator (Fig. 3, Supplementary Table S3). Edugator was isolated from soil using *Mycobacterium smegmatis* as a host through the SEA-PHAGES program (43); PhagesDB). However, this annotation does not establish Edugator—or even a phage antigen— as the source of the immune response. Scaffolding proteins mediate capsid assembly but are typically absent from mature virions. Recognition of this oligopeptide could therefore reflect exposure to scaffolding proteins released during phage replication and bacterial lysis or cross-reactivity with homologous sequences from other phages or organisms. Current sequence databases do not allow a potential non-phage immunogen to be identified (Fig. 5, Supplementary File 2); resolving its origin will require epitope-level analysis and experimental validation of candidate homologous antigens.

PhageScan, like other PhIP-Seq approaches, has several limitations. Because it uses short peptides, it primarily detects linear epitopes and may fail to capture conformational or discontinuous epitopes, as well as those dependent on post-translational modifications. Uneven representation of phage proteomes and variability in phage display efficiency may also bias epitope coverage and detection sensitivity. Moreover, sequence enrichment provides a relative measure of antibody binding rather than a direct estimate of antibody concentration, titre, or affinity. The assay measures binding but does not determine whether the detected antibodies neutralize phages, affect protein function, or confer protection; consequently, some signals may be nonspecific or biologically irrelevant. Finally, PhageScan characterizes circulating humoral responses but does not capture cellular immunity or tissue- and mucosa-specific antibody repertoires (44,45).

Despite these limitations, PhageScan provides a highly multiplexed and scalable approach for profiling antibody reactivity against hundreds of thousands of phage-derived sequences using small serum volumes. Unlike whole-protein assays, it maps antibody recognition at oligopeptide-level resolution, enabling reactive regions to be assigned to specific proteins and functional domains. This resolution facilitates the identification of shared and individualized antibody profiles, potential cross-reactive epitopes, and candidate sequences with functional or therapeutic relevance. PhageScan is therefore particularly valuable as a discovery and hypothesis-generating platform, with selected findings subsequently validated using complementary serological and functional assays (46,47).

Accordingly, PhageScan may have broad applicability in studies of phage-antibody interactions, particularly for investigating the development of epitope-specific responses, cross-reactivity among phages, and the immunological regulation of phage persistence. Because phage-specific IgG may constrain phage survival (48), antibody-mediated selection could shape phageome composition and thereby indirectly influence bacterial communities and broader microbiome dynamics. These insights may also inform phage therapy, including treatment planning and the development of ready-to-use therapeutic phage collections, as well as the selection of phages for drug- and vaccine-delivery platforms.

## Supporting information

Supplementary File 1 (Interactive HTML visualization)

Supplementary File 2 (Interactive HTML visualization)

## Acknowledgements

This work was supported by the National Science Centre in Poland: grant no. 2019/35/B/NZ7/01824. The authors thank Anna Wilk (StatSoft Polska Sp. z o.o., Kraków, Poland) for her valuable contributions to the statistical analyses and providing expert advice.

## Supplementary

Supplementary Table S1. Curation and categorization of GenBank metadata for phage isolation sources and bacterial hosts.

Supplementary Table S2. Library-normalized measure of virus-specific IgG recognition (burden score) calculated for phage species.

Supplementary Table S3. Complete list of immunogenic phage-derived oligopeptides identified in this study

Supplementary Table S4. Representative sequence similarities between immunogenic phage-derived oligopeptides and sequences from other phages or bacteria.

Supplementary Table S5. Phage-derived oligopeptides identified as immunogenic separately in the Polish and US cohorts.

Supplementary Table S6. Comparison of normalized PhageScan signals for the ImmE epitope with ImmE-specific ELISA reactivity, all available samples.

Supplementary File 1. Identification of amino acid residues involved in antigen recognition within the most frequently recognized phage-derived oligopeptides by human IgG. This file contains an interactive visualization in HTML format that should be downloaded, saved, and opened in a web browser.

Supplementary File 2. Analysis of sequence homology between recognized phage-derived oligopeptides and eukaryotic organisms. This file contains an interactive visualization in HTML format that should be downloaded, saved, and opened in a web browser

Supplementary File 3. Optimization of ImmE activity in Sytox assay on Lactococcus (PDF)

## Suplementary File 3

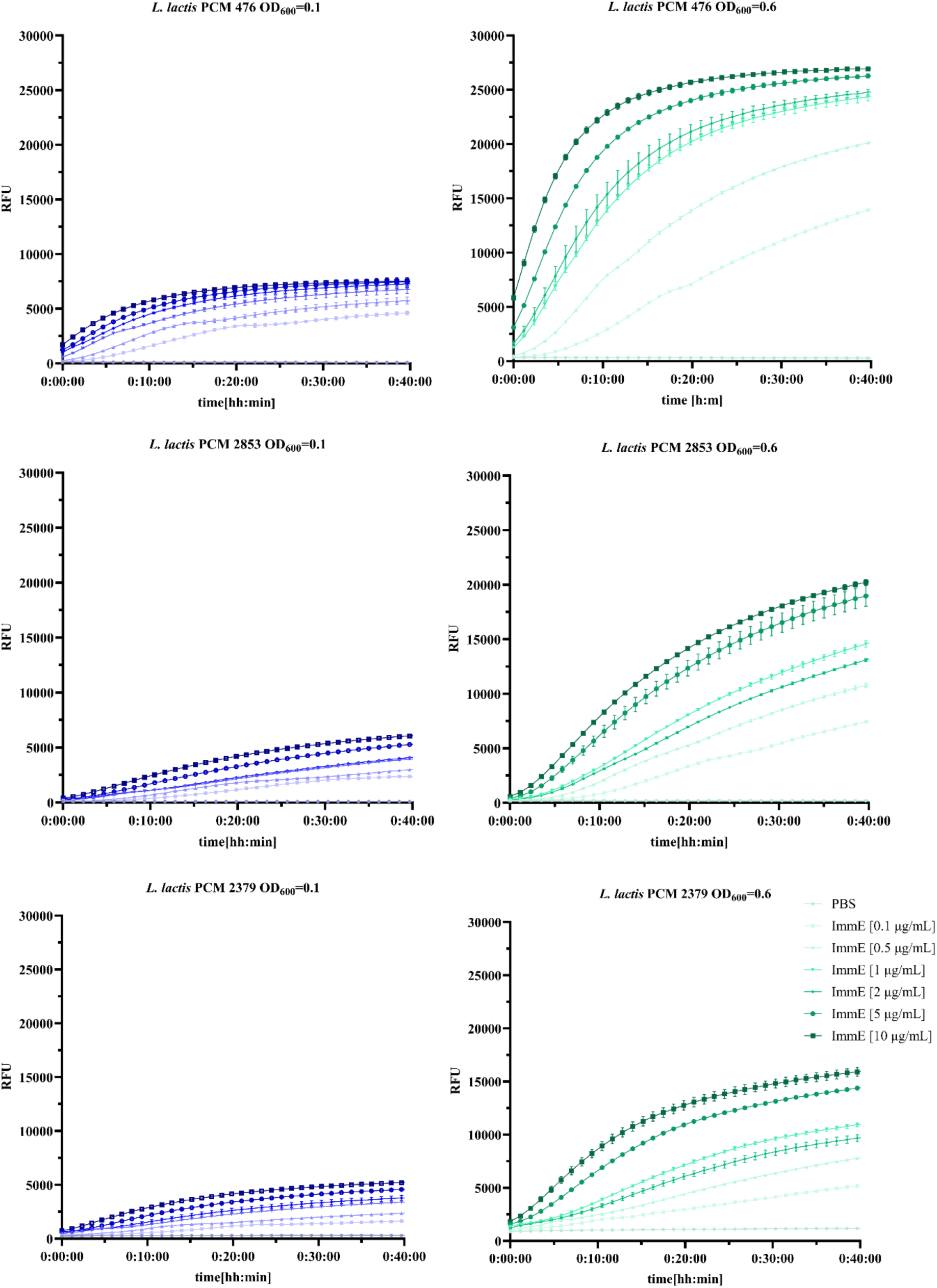

## Notes

### Competing Interest Statement

The authors have declared no competing interest.

