## Supplementary File 1 (Interactive HTML visualization) for "A synthetic phageome reveals widespread and diverse human antibody responses to bacteriophages": Supplementary File 1.html

Oligopeptide Responders


Search: 
Filter:
All
Has red
Has yellow
No red
Only green
**Legend:**adj.p < 0.00050.0005 ≤ adj.p ≤ 0.05adj.p > 0.05

| Responders (% of cohort) | Oligopeptide | Sequence |
| --- | --- | --- |
