## Supplementary File 2 (Interactive HTML visualization) for "A synthetic phageome reveals widespread and diverse human antibody responses to bacteriophages": Supplementary File 2.html

BLAST Protein Homology — Dashboard


### BLAST Protein Homology — Dashboard

Generated from output Excel

12,576

Total input

1,359

Significant

11,217

Not significant

The pie chart displays the proportional distribution of query sequences classified as
significant or not significant based on BLAST/DIAMOND homology search results against the
eukaryotic reference database. A sequence is classified as significant if at least one of
its reported hits carries an E-value ≤ 0.05, indicating a statistically meaningful
alignment. The inner ring (donut format) shows absolute sequence counts for each category,
while the percentage values reflect each category's share of the total dataset. A high
proportion of significant sequences suggests widespread conservation of the queried proteins
among eukaryotic organisms.

Click a bar to see organism breakdown

The grouped bar chart illustrates the distribution of the number of eukaryotic homologous
hits identified per query sequence, ranging from 0 (no hits found) to 5 (the maximum number
of hits reported per sequence). Each hit-count category is shown as a pair of bars:
**green** represents sequences classified as
significant (at least one hit with E-value ≤ 0.05), and
**gray** represents sequences that are not significant.
Both counts are printed above their respective bars, so every value stays legible
regardless of bar height. Sequences with zero hits carry no database match at all and are
therefore always not significant. Clicking any bar reveals a drill-down table listing the eukaryotic
organisms most frequently represented among the hits in that group, enabling rapid
assessment of taxonomic diversity across hit-count categories.

Click a bar to see organism breakdown

The histogram presents the distribution of best-hit E-values for sequences classified as
significant. The E-value (Expect value) is a statistical parameter reflecting the number of
alignments with an equivalent or better score expected by chance alone in a database of the
given size; lower E-values therefore indicate stronger, more statistically significant
matches. Bars are color-coded using a blue gradient — dark blue corresponds to very low
E-values (strongest, most significant hits) and light blue to E-values approaching the
0.05 significance threshold (weakest qualifying hits). Only sequences flagged as significant
are included in this chart. Clicking a bar displays the breakdown of eukaryotic organisms
contributing hits within that E-value interval.

The scatter plot visualises the relationship between hit significance and sequence
similarity for the best hit of each significant query sequence. The x-axis represents
−log₁₀(E-value), such that higher values correspond to more statistically
significant alignments. The y-axis shows the percentage of identical amino acid residues
within the aligned region (% identity). Each point is colored by % identity according to the
Viridis colorscale (dark purple = low identity, bright yellow = high identity).
Hovering over a point reveals the protein name, matched organism and raw E-value. Sequences
clustering in the upper-right region represent strong, highly conserved homologs, whereas
points in the lower-left indicate more distant matches.

Click a bar to see organism breakdown

The histogram shows the distribution of percent sequence identity values for the best hit
of each significant query sequence. Identity percentage reflects the fraction of aligned
positions at which the query and matched database sequences share an identical amino acid
residue. Higher identity values suggest closer evolutionary relationships between the query
protein and its eukaryotic homolog. The shape of the distribution provides insight into the
overall degree of conservation across the dataset: a unimodal peak in the high-identity
range is characteristic of well-conserved protein families, while a broad or left-skewed
distribution indicates heterogeneous levels of homology. Clicking a bar reveals the
organism breakdown for that identity interval.

Click a bar to see organism breakdown

The histogram presents the distribution of query coverage values for the best hit of each
significant sequence. Query coverage represents the percentage of the query sequence length
that participates in the alignment with the matched database entry. High coverage
(80–100%) indicates that nearly the entire query protein was matched, suggesting
structural and functional conservation across its full length, while low coverage may
indicate partial domain homology, the presence of lineage-specific insertions, or
incomplete alignment. Only significant sequences are included. Clicking any bar displays a
breakdown of the eukaryotic organisms represented within that coverage range.

The horizontal bar chart lists the 20 most frequently occurring eukaryotic organisms across
all hits reported for significant query sequences. The x-axis denotes the total number of
individual hit records attributed to each organism, aggregated across all five hit positions
(hit 1–hit 5). Organisms with the longest bars are the most common sources
of homologous sequences in the eukaryotic reference database relative to the query dataset,
and may represent phylogenetically close relatives or highly sequenced model organisms. The
chart provides an overview of the overall taxonomic composition of the identified eukaryotic
homologs and can highlight potential biases in database representation.

The horizontal bar chart presents the top cluster ancestor accessions from the clustered NR
database (ClusteredNR), aggregated across all hits of significant sequences. In clustered
BLAST searches, homologous sequences are grouped into clusters, each represented by a
canonical sequence and associated with a cluster ancestor that reflects the inferred
evolutionary root of that cluster. Higher hit counts for a given ancestor indicate that the
corresponding sequence cluster contributed more frequently as a source of eukaryotic
homologs across the query dataset. This chart is populated only when the NCBI BLAST engine
is used; results generated with the DIAMOND local alignment tool may not include cluster
ancestor information.

Close
